# Identification of anti-norovirus genes in mouse and human cells using genome-wide CRISPR activation screening

**DOI:** 10.1101/350090

**Authors:** Robert C. Orchard, Meagan E. Sullender, Bria F. Dunlap, Dale R. Balce, John G. Doench, Herbert W. Virgin

## Abstract

Noroviruses (NoVs) are a leading cause of gastroenteritis world-wide, yet host factors that restrict NoV replication are not well understood. Here, we use a CRISPR activation (CRISPRa) genome-wide screening to identify host genes that can inhibit murine norovirus (MNoV) replication in either mouse or human cells. Our screens identified with high confidence 57 genes that can inhibit MNoV infection when overexpressed. A significant number of these genes are in interferon and immune regulation signaling networks, but surprising, the majority of the genes identified are not associated with innate or adaptive immunity nor with any antiviral activity. Confirmatory studies of eight of the genes in validate the initial screening data. Mechanistic studies on *TRIM7* demonstrated a conserved role of the molecule in mouse and human cells in restricting MNoV in a step of infection after viral entry. Furthermore, we demonstrate that two isoforms of *TRIM7* have differential antiviral activity. Taken together these data provide a resource for understanding norovirus biology and demonstrate a robust methodology for identifying new antiviral molecules across cell types and species.

**Author Summary:** Norovirus is one of the leading causes of foodborne illness world-wide. Despite its prevalence, our understanding of norovirus biology is limited due to the difficulty in growing human norovirus *in vitro* and a lack of an animal model. Murine norovirus (MNoV) is a model norovirus system because MNoV replicates robustly in cell culture and in mice. To identify host genes that can restrict norovirus replication when overexpressed we performed genome-wide CRISPR activation (CRISPRa) screens to induce gene overexpression at the native locus through recruitment of transcriptional activators to individual gene promoters. We found 57 genes could block murine norovirus replication in either mouse or human cells. Several of these genes are associated with classical immune signaling pathways, while many of the molecules we identified have not been previously associated with antiviral activity. Our data is a resource for those studying norovirus and we provide a robust approach to identify novel antiviral genes.

## Introduction

Noroviruses are non-enveloped viruses with positive-sense RNA genomes that infect the gastrointestinal tract of mammals [1]. Human norovirus (HNoV) is the leading cause of viral gastroenteritis worldwide and is estimated to have an economic burden of $1 billion annually [2]. Despite the high infectivity of HNoV between individuals, it has proven difficult to culture *in vitro* with significant advances made only in the past few years [3, 4]. Because of the challenges in growing HNoV in cell culture and lack of an animal model, murine norovirus (MNoV) has emerged as a system for understanding norovirus biology. MNoV grows robustly in cell culture, is amenable to reverse genetics, and is a natural pathogen of laboratory mice. MNoV is similar to HNoV in that both viruses are spread fecal-orally, infect intestinal tissues, have similar genomic organization and protein function, and can establish persistent infection in individuals [1].

Host factors that limit or modulate norovirus replication are still not well understood. We and others previously used a whole-genome CRISPR/Cas9 loss-of-function screen to identify essential host factors required for MNoV growth in macrophage-like cell lines leading to the identification of CD300lf as a proteinaceous receptor for MNoV [5, 6]. Importantly, expression of mouse CD300lf in cells is sufficient for replication in all cell types tested including human cells, thus expanding the repertoire of cell lines that can be utilized to study norovirus infection with MNoV [5, 6]. These studies also demonstrate that the cellular machinery for replication of MNoV is highly conserved with species specificity conferred by the receptor. While CRISPR knockout studies for MNoV and other viruses have identified host factors necessary for viral infection, they have yet to identify restriction factors that antagonize viral replication [5–11]. Recent modifications of the CRISPR/Cas9 system has led to the ability to transactivate gene expression through specific recruitment of transcriptional activators to individual promoters and has been broadly called CRISPR activation (CRISPRa; [12–14]). cDNA overexpression systems have been used to identify viral restriction factors but this approach has several technical hurdles, most notably the cloning, delivery, and expression of large transcripts and the need to select individual protein isoforms rather than screening for the effects of all transcripts that emanate from a given cellular gene promoter. Because CRISPRa induces gene expression from the native locus and relies only on sgRNA homology for specificity, it overcomes many of these obstacles. We therefore utilized CRISPRa to identify genes whose overexpression block MNoV replication.

We use recently developed CRISPRa genome-wide libraries to screen for host genes that antagonize MNoV replication in both mouse and human cells. Our screens and subsequent validation identify numerous anti-MNoV genes including canonical interferon stimulated genes and several genes with novel antiviral activity. We demonstrate that the antiviral activity of one such gene, *TRIM7*, is highly specific to only one of the two major *TRIM7* isoforms, highlighting the utility and power of the CRISPRa screening approach. Further mechanistic studies demonstrate that TRIM7 inhibits MNoV replication post viral entry. These data provide a resource for understanding norovirus biology and a methodology for identifying unappreciated viral restriction factors.

## Results

### Generation of CRISPRa libraries in mouse and human cells

We set out to identify genes in mouse and human cells that have anti-norovirus activity when overexpressed. By screening cell lines from two different species, we sought to capture a wider diversity of antiviral genes and potentially identify those molecules with evolutionary conserved antiviral function. We first generated cell lines that can be infected by MNoV and express the CRISPRa machinery. For the mouse screen, we used BV2 microglial cells, previously used for loss-of-function CRISPR screening, in which MNoV replicates robustly [5]. As noroviruses exhibit strict species tropism and at this time HNoV replication is not robust or scalable for whole-genomic pooled screens, we selected HeLa cells to use as a model human cell line and generated HeLa-CD300lf stable cells that can be infected by MNoV. We introduced the catalytically dead Cas9-VP64 (dCas9-VP64) fusion construct into both the BV2 and HeLa-CD300lf cell lines (BV2-dCas9-VP64 and HeLa-CD300lf-dCas9-VP64).

Before proceeding with a genome-wide screen, we developed a simple assay to test transcriptional activation activity in BV2-dCas9-VP64 and HeLa-CD300lf-dCas9-VP64 cells relying on the low level of expression of CD4 in these cells. sgRNAs targeting the CD4 transcriptional start site robustly increased the surface levels of CD4 in BV2-dCas9-VP64 and HeLa-CD300lf-dCas9-VP64 cells compared to an empty vector control (Figure 1A-1B).

We next introduced a pooled, genome-wide CRISPRa library into BV2-dCas9-VP64 (Caprano library) and HeLa-CD300lf-dCas9-VP64 (Calabrese library) cells. Importantly, these libraries use an updated CRISPRa sgRNA vector system (Sanson et al. in preparation). This system (pXPR_502) is bicistronic with one promoter expressing the sgRNAs with PP7 stem loops while the second promoter drives expression of the synergistic transcriptional activators (Figure 1A). This new system is advantageous over the previous synergistic activation mediator (SAM) vectors as it consolidates the transcription activators and sgRNA into one lentivirus [14]. Caprano and Calabrese libraries contain 134,076 and 113,240 total sgRNAs, respectively, with each gene targeted with 3-6 sgRNAs split between two pools, SetA and SetB. After introduction of these libraries into BV2-dCas9-VP64 and HeLa-CD300lf-dCas9-VP64 cells, we challenged the resulting pools with either MNoV^CW3^ and MNoV^CR6^ and isolated resistant cells (Figure 1D). MNoV^CW3^ causes acute, systemic infection in mice while MNoV^CR6^ establishes a persistent infection in intestinal tuft cells, a specialized intestinal epithelial cell [15–17]. MNoV^CW3^ and MNoV^CR6^ have been used to model human noroviruses, transkingdom interactions, and innate immunity restriction of viral tropism [4, 18–23]. In summary, we performed four independent CRISPRa screens using two different cell lines (BV2 vs. HeLa) and two different viral strains (MNoV^CW3^ and MNoV^CR6^) in an effort to provide a broad resource for understanding norovirus biology in multiple cellular settings.

### CRISPRa screen for anti-MNoV in BV2 cells

BV2-dCas9-VP64 cellular pools containing the Caprano library were infected with MNoV^CW3^ or MNoV^CR6^ at an MOI of 5. Surviving cells were harvested after a brief expansion period (Figure 1D). sgRNAs from resistant cells were compared to mock infected control using STARS (Doench et al., 2016). The STARS algorithm calculates a score and a false discovery rate (FDR) for a gene with at least two sgRNAs enriched in the top 10% of a ranked ordered list of total sequenced sgRNAs. We consider a gene a hit in an individual screen if the FDR was < 0.25. Five and seven genes met this criterion in BV2 cells that survived MNoV^CW3^ or MNoV^CR6^ infection, respectively. Four genes (*Trim7*, *Prdm1*, *Tbx19*, and *Obox6*) were enriched after challenge with both viral strains (Tables S1-S2 and Figure 2A-2B). Gene Set Enrichment Analysis (GSEA) of the screen hits demonstrated an enrichment in interferon gamma and alpha response pathways, consistent with the known role of these responses to limit viral replication (Table 1). Overall these results demonstrate the ability of the CRISPRa technology in identifying antiviral activity of overexpressed genes.

**Table 1:** GSEA of BV2 CRISPRa screening hits.

| Gene Set Name | # Genes in Overlap (k) | p-value | FDR q-value |
| --- | --- | --- | --- |
| GO RESPONSE TO INTERFERON BETA | 3 | 2.71E-08 | 1.62E-04 |
| GO NEGATIVE REGULATION OF MULTI ORGANISM PROCESS | 4 | 7.83E-08 | 2.33E-04 |
| GO NEGATIVE REGULATION OF VIRAL GENOME REPLICATION | 3 | 3.23E-07 | 6.43E-04 |
| GO REGULATION OF VIRAL GENOME REPLICATION | 3 | 1.18E-06 | 1.76E-03 |
| GO NEGATIVE REGULATION OF VIRAL PROCESS | 3 | 1.98E-06 | 2.36E-03 |
| HALLMARK INTERFERON ALPHA RESPONSE | 3 | 2.57E-06 | 2.55E-03 |
| GO REGULATION OF MULTI ORGANISM PROCESS | 4 | 7.18E-06 | 6.12E-03 |
| GO DEFENSE RESPONSE TO VIRUS | 3 | 1.24E-05 | 8.35E-03 |
| GO NEGATIVE REGULATION OF VIRAL ENTRY INTO HOST CELL | 2 | 1.26E-05 | 8.35E-03 |
| GO RESPONSE TO INTERFERON ALPHA | 2 | 1.40E-05 | 8.35E-03 |
| HALLMARK INTERFERON GAMMA RESPONSE | 3 | 2.25E-05 | 1.19E-02 |
| GO REGULATION OF SYMBIOSIS ENCOMPASSING MUTUALISM THROUGH PARASITISM | 3 | 2.42E-05 | 1.19E-02 |
| GO PHOSPHATIDYLSERINE METABOLIC PROCESS | 2 | 2.78E-05 | 1.19E-02 |
| GO REGULATION OF VIRAL ENTRY INTO HOST CELL | 2 | 2.78E-05 | 1.19E-02 |
| GO CELLULAR MODIFIED AMINO ACID METABOLIC PROCESS | 3 | 2.99E-05 | 1.19E-02 |
| GO RESPONSE TO VIRUS | 3 | 4.22E-05 | 1.53E-02 |
| GO ALDITOL PHOSPHATE METABOLIC PROCESS | 2 | 4.37E-05 | 1.53E-02 |
| GO CELLULAR MODIFIED AMINO ACID BIOSYNTHETIC PROCESS | 2 | 1.01E-04 | 3.35E-02 |

### CRISPRa screen for anti-MNoV genes in HeLa-CD300lf cells

In comparison to BV2 cells, MNoV exhibits less selective pressure in HeLa-CD300lf cells and thus an MOI of 50 for MNoV^CW3^ and MNoV^CR6^ was used, additionally, a second challenge of MNoV^CR6^ was preformed to exert a high selective pressure for this strain of virus (see methods for more details). sgRNAs were sequenced from resistant cells and compared to a mock infected control using the STARS algorithm [24]. 33 and 22 genes in MNoV^CW3^ and MNoV^CR6^ infected HeLa-CD300lf, respectively, scored as hits (Tables S3-S4 and Figure 2A and 2C). Six genes (*TRIM7*, *PITX1*, *HOXC11*, *DDX60*, *MX1*, and *PLSCR1*) were enriched after challenge with both viruses.

To explore the possibility that individual hits function in a similar pathway, we took the 49 genes that scored for either viral strains and analyzed them using GSEA [25, 26]. Comparison of the hallmark genes with our dataset indicated a significant enrichment in genes related to interferon alpha responses, including *GBP2*, which has been recently shown to block MNoV replication complex formation [27]. When analyzing the function of the genes by GO terms we also found an enrichment for interferon and cytokine responses along with genes involved in embryogenesis and RNA polymerase gene expression (Table 2). These data demonstrate the ability to discover expected candidate antiviral genes (e.g. those regulated by inflammatory cytokines) and candidate genes involved in other pathways (e.g. embryogenesis).

**Table 2:**
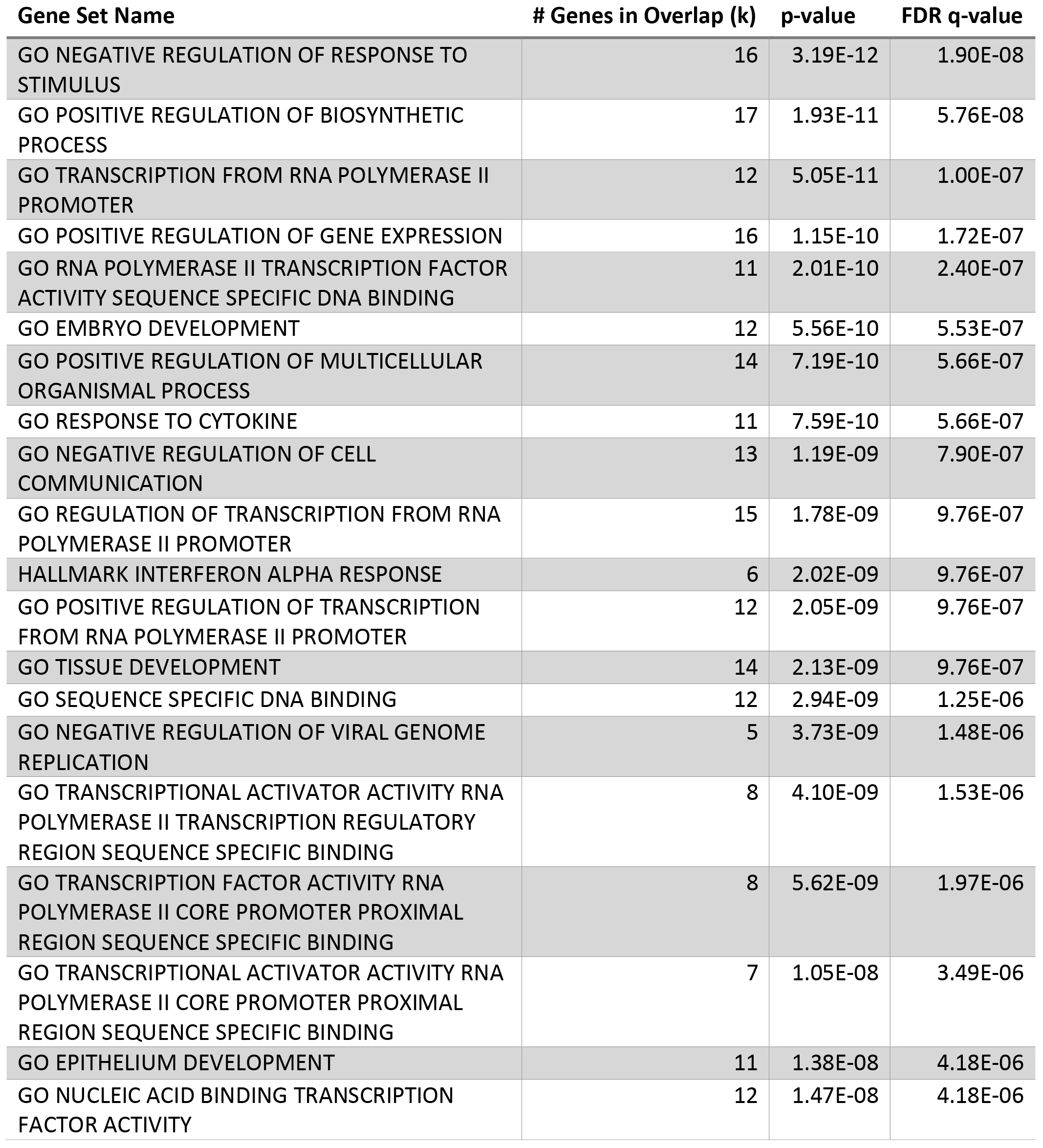
GSEA of HeLa-CD300lf CRISPRa screening hits.

### Analysis and comparison of screen in BV2 and HeLa cells

We compared the results from the BV2 and HeLa-CD300lf CRISPRa screens. The top hit in all virus and cell type combinations was *TRIM7* (Figure 2A and Tables S1-S4). No other gene had an FDR <0.25 and only *TRIM7* and *PLSCR1* had a STARS score > 2.0 in all four screens (Figure 2A). Similar results are obtained when we subdivide the screens based upon viral strain (Tables S1-S4). When looking at the GSEA both screens had an interferon signature (Table 1 and 2). The lack of concordance between other hits could be due to cell type, species, library, or selective pressure differences between screens. Additionally, we find that false-negatives are common (see below) and our data likely underrepresents the true overlap of function across the cellular systems. None the less, these data highlight the importance of examining multiple cell types for antiviral genes.

### Validation of CRISPRa screening results in HeLa cells

We validated our screening results in human cells. We individually, introduced 2 sgRNAs per gene into HeLa-CD300lf-dCas9-VP64 cells targeting *TRIM7*, *MUC1*, *LRRC15*, *PLSCR1*, *HOXC11*, *GBP2*, *MX1*, *BLNK* and compared to cells transduced with an empty pXPR_502 vector or pXPR_502 with a sgRNA targeting CD45. These genes encompass seven of the top 15 enriched genes after MNoV^CW3^ challenge. Additionally, three of the eight genes showed enrichment in an MNoV-strain dependent manner (MUC1 and GBP2 for MNoV^CW3^ and BLNK for MNoV^CR6^) and thus were ideal for testing strain specificity for potentially antiviral molecules.

As the screen was designed to enrich for genes that protected cells from virus induced cytopathic effect (CPE), we first tested the ability of cells to survive MNoV infection. All genes tested had at least one sgRNA show statistical growth advantage over control cells when challenged with either MNoV^CW3^ or MNoV^CR6^ (Figure 3A and 3B). Furthermore, four of the eight and five of the eight genes had both sgRNAs provide a survival phenotype when challenged with MNoV^CW3^ and MNoV^CR6^, respectively (Figure 3A and 3B). We were unable to validate an MNoV strain-specific protective effect for any gene (Figure 3C). Taken together, these data validate the reproducibility of the CRISPRa screening platform and indicate the potential for false-negatives.

We next tested whether overexpression of these genes inhibited production of infectious virions by analyzing the amount of MNoV replication 24 hours-post infection (Figure 3D and 3E). Six of the eight (MNoV^CW3^) five of the eight (MNoV^CR6^) genes tested had at least one sgRNA significantly reduce the amount of MNoV produced (Figure 3D-3F). Additionally, both sgRNAs for three genes (*TRIM7*, *GBP2*, and *MX1*) blocked the production of MNoV^CW3^ and MNoV^CR6^. In summary these data provide the first systematic overview of the molecules that can inhibit MNoV infection in human cells.

### Isoform specificity of the antiviral activity of TRIM7

While we have identified a significant number of novel antiviral molecules, our overarching goal is to identify molecules that broadly inhibit NoV replication in multiple cellular contexts. Given that the top gene in all four screens was *TRIM7*, we decided to focus our efforts on characterizing the anti-MNoV activity of *TRIM7* in HeLa and BV2 cells. TRIM7 (also GNIP) is an E3-ligase previously shown to interact with glycogenin and ubiquitinate the AP-1 coactivator RACO1; however the role of TRIM7 during viral infection has not been explored [28, 29].

TRIM7 has at least 4 isoforms in humans with isoform 1 and isoform 4 being the most commonly studied and display different expression pattern in muscular tissues and cells [30]. We generated stable BV2 and HeLa-CD300lf cells overexpressing human TRIM7 isoform 1 or isoform 4 (Figure 4A). While expression of TRIM7 isoform 1 protected both HeLa-CD300lf and BV2 cells from MNoV CPE, TRIM7 isoform 4 provided no protection (Figure 4B-4E). TRIM7 isoform 1, but not isoform 4, reduced the growth of MNoV^CW3^ and MNoV^CR6^ in both human and mouse cells (Figure 4F-4I). Taken together, these data demonstrate an isoform specificity for the anti-MNoV mechanism that is conserved across species.

Finally, we determined which step of the viral life cycle was targeted by TRIM7. Since ubiquitination can lead to proteaseome-mediated degradation of proteins, we first tested whether the MNoV receptor, CD300lf, was reduced upon TRIM7 overexpression. In HeLa-CD300lf we found no effect of TRIM7 on CD300lf protein levels (Figure 4A). We next tested the hypothesis that TRIM7 targets a critical step of MNoV entry. Transfection of MNoV^CW3^ RNA was unable to rescue MNoV production in human TRIM7 isoform 1 expressing BV2 cells indicating that TRIM7 can target a post-entry step of the MNoV life cycle to inhibit viral replication (Figure 4J).

## Discussion

Here we present a genome-wide screen to identify genes that can inhibit murine norovirus replication upon overexpression in both mouse and human cells. Our results provide a resource for those studying norovirus biology and validate CRISPRa screening as an approach to uncover novel antiviral molecules. As human norovirus studies are limited due to the difficulty in culturing the virus in cell culture, we speculate that our screens with MNoV in mouse and human cells may identify evolutionary conserved genes that may broadly inhibit NoV replication. Such genes might be targeted through genetic or chemical perturbagens to enhance HNoV replication in cell culture systems, or to generate therapeutic effects.

With the discovery that Cas9 binding to DNA does not require nuclease activity, several new technologies have emerged to synthetically modulate mammalian genomes with unprecedented precision and scale [31]. By leveraging the cytotoxicity of lytic viruses, these pooled screening approaches are ideal to identify genes that alter viral infection. While most investigations using CRISPR/Cas9 genome-wide screens have focused on determining host factors that facilitate viral infection, our results highlight the utility of CRISPRa to identify antiviral molecules.

Genes identified in these screens exhibited enrichment in viral and immune related genes not previously appreciated to have anti-MNoV activity (Figure 2A). Given that the tropism of both acute and persistent strains of MNoV are dictated by interferons, these data provide the first insight into potential interferon stimulated genes (ISGs) that mediate viral restriction of MNoV [18, 19, 21]. It will be important to determine whether these ISGs contribute to determining norovirus tropism *in vivo.*

Surprisingly, in addition to the ISGs, a number of genes not appreciated to have immune functions were identified as possessing anti-MNoV activity (Figure 2A). These represent unique opportunities to identify novel norovirus vulnerabilities and to determine whether these genes possess antiviral activity against other viruses that may explain tropism or be exploited for therapeutic potential. One of these genes, TRIM7, inhibited MNoV replication after viral entry (Figure 4I). As ubiquitination can have diverse effects on protein stability and function, it will be essential to determine the substrate and type of lysine-linkage in order to understand the mechanism of viral restriction [32]. Additionally, we mapped the antiviral function of TRIM7 to a specific isoform, while a closely related isoform had no detectable anti-MNoV activity (Figure 4A-4H). The only differences between these isoforms are within the protein-protein interaction coiled-coil domain SPRY domains. As these molecules can be differentially and coexpressed in different cellular contexts and TRIM proteins are capable of homodimerization, it will be important to determine if isoform 4 can regulate the antiviral activity or the normal physiological functions of isoform1 [33]. Furthermore, as traditional cDNA based screens require the scientist to select a specific isoform, CRISPRa approaches enable complex transcriptional variance to be reduced and encompassed by sgRNAs, which is highlighted by the specificity of anti-MNoV by TRIM7 isoforms [14].

In conclusion, our data provides the first systematic insight into genes that can restrict norovirus replication when overexpressed and provides a methodological approach to identify novel antiviral molecules using CRISPRa technology that may be broadly applicable to other viral systems.

## Methods

### Cell culture

BV2 cells (BV-2 cells were provided by Dr. Yuanan Lu, University of Hawaii at Manoa), 293T (ATCC) and Hela cells (ATCC) were cultured in Dulbecco’s Modified Eagle Medium (DMEM) with 10% fetal bovine serum (FBS), and 1% HEPES. HeLa cells were transduced with pCDH-MSCV-CD300lf-T2A-Hygromycin and selected with Hygromycin for 2 weeks prior to sorting CD300lf high expressing cells to generate HeLa-CD300lf-Hygromycin cells. For BV2 cells, 4 μg/ml of puromycin (Sigma Aldrich) and 5 μg/ml blasticidin (Invitrogen), was added where appropriate. For HeLa cells, 1 μg/ml of puromycin, 5 μg/ml blasticidin, and 350 μg/ml hygromycin (Invitrogen) were added as appropriate.

Lentivirus was generated by transfecting lentiviral vectors with packaging vector (psPAX2) and pseudo-typing vector (pCMV-VSV-G) into 293Ts using TransIT-LT1 (Mirus). 48 hours post-transfection, supernatants were collected, filtered through a 0.45 μm filter (Millipore), and added to the indicated cells. After 48 hours, cells were selected with the appropriate antibiotic.

### Plasmids

Guide constructs were cloned into pxpr_502 (Addgene #96923). A complete list of guide sequences used in this study can be found in the supplemental materials. Human *TRIM7* isoform 1 (NP_976038.1) and isoform 4 (NP_203128.1) were codon optimized and cloned into pCDH-MCS-T2A-Puro-MSCV (Systems Biosciences #CD522A-1). Codon optimized cDNA sequences for the constructs can be found in the supplemental materials. All plasmids were sequenced verified prior to use.

### Transcriptional Activation Reporter Assays

BV2 cells and HeLa-CD300lf cells were transduced with pLenti-dCas9-VP64-Blast [14]. Transcriptional activity was assessed in these cells by transducing either parental or dCas9-VP64 cells with pxpr_502 lentiviruses targeting CD4. Cells were transduced for 2 days and subsequently, selected for 7 days with puromycin and the frequency of CD4 expression was assessed by flow cytometry.

### BV2 CRISPRa screen

BV2-dCas9-VP64 cells were transduced with Set A and SetB of the caprano CRISPRa library at an MOI of 0.3 to 0.5 (Sanson et al. in preparation). Transductions were performed with enough cells to enable each sgRNA in the library to be represented at least 500 times. Each experimental condition was performed in duplicate with each replicate consisting of 3.0 × 10^7^ BV2 cells expressing the complete CRISPRa system seeded equally across three 150cm2 dishes on day five posttransduction. Cells were infected 24 hours later with either MNoV^CW3^ or MNoV^CR6^ at MOI of 5. Mock infected cells were harvested 48 hours after seeding. Ten days postinfection surviving cells were harvested. Genomic DNA was isolated from cells using Mini (<5 × 10^6^ cells), Midi (5 × 10^6^ - 3 × 10^7^ cells), and Maxi (3 × 10^7^ cells - 1 × 10^8^ cells), kits (Qiagen) according to the Broad Institute’s Genetic Perturbation Platform protocols (https://portals.broadinstitute.org/gpp/public/resources/protocols).

### HeLa CRISPRa screen

HeLa-CD300lf cells expressing dCas9-VP64 were transduced with the Set A and SetB of the caprano CRISPRa library at an MOI of 0.3. Each Calabrese pool was delivered by lentiviral transduction of 1.2 × 10^8^ HeLa cells at an MOI ~0.3. This equates to 3.6 × 10^7^ transduced cells, which is sufficient for the integration of each sgRNA at least 500 independent times. Two days post transduction, puromycin was added to the media and transduced cells were selected for a total of 7 days. 5 × 10^6^ HeLa CRISPRa cells were seeded in a T175 tissue culture flask and for each experimental conditional was done in duplicate with 6 independent flasks per replicate (3e7 cells per replicate). Cells were infected with either MNoV^CW3^ or MNoV^CR6^ at an MOI 50 at the time of seeding. Two days after infection, mock infected cells were harvested for genomic DNA extraction. At this time point, roughly 85% of the MNoV^CW3^ infected cells displayed cytopathic effect while MNoV^CR6^ cells had 35-50% and both sets of conditions had cells washed twice with PBS and fresh media added. After an additional 2 days of viral challenge MNoV^CR6^ challenged cells had roughly 25% viable cells and thus were washed, trypsinized, collected and counted and replated at 5 × 10^6^ cells/flask and rechallenged. The MNoV^CW3^ challenged cells had <10% viable at this time point and thus only a single challenge was performed. Cells were harvested after the second challenge of MNoV^CR6^ and the single challenge of MNoV^CW3^ when they started to form discernable, stable colonies, between 7-14 days from the initial challenge. All flasks from a single replicate were harvested together. Genomic DNA was isolated from surviving cells using a Qiamp DNA mini kit according to manufacturer instructions (Qiagen).

### CRISPR screen sequencing and analysis

Illumina sequencing and STARS analysis was performed as described previously [5]. Briefly, genomic DNA was aliquoted into multiple wells of a 96-well plate with up to 10 μg of DNA in 50 μl total volume. A PCR mastermix consisting of ExTaq DNA polymerase (Clontech), ExTaq buffer, dNTP, P5 stagger primer, and water was generated. 40 μl of PCR master mix, and 10 μl of a barcoded primer was added to each well containing 50 μl of DNA. Samples were PCR amplified as follows: 95°C for 1min., followed by 28 cycles of 94°C for 30 sec., 52.5°C for 30sec., 72°C for 30 sec. with a final 10 min. at 72°C. PCR product was purified with Agencourt AMPure XP SPRI beads according to the manufacture’s protocol (Beckman Coluter). Samples were sequenced on an Illumina HiSeq 2000. Barcodes in the P7 primer were deconvuluted and the sgRNA sequence was mapped to a reference file of sgRNAs in the caprano or calabrese library. To normalize for different numbers of reads per condition, read counts per sgRNA were normalized to 10^7^ total reads per sample. This normalized value was then log-2 transformed. sgRNAs that were not sequenced were arbitrarily assigned a read count of 1. Replicates were then averaged together and sgRNA frequencies were then analyzed using STARS software, which is available at http://www.broadinstitute.org/rnai/public/software/index [24]. STARS computes a score for each gene of rank-ordered sgRNA hits that was above 10% of total sequenced sgRNAs. A STAR score was assigned to genes that had a sgRNA score above this threshold in either of the two independent pools.

For GSEA analysis, screen hits were queried at the BROAD GSEA portal (https://software.broadinstitute.org/gsea/index.jsp). Only the top 20 gene set names with an FDR < 0.05 are shown in the tables.

### MNoV Assays

MNoV^CW3^ (Gen bank accession no. EF014462.1) and MNoV^CR6^ (Gen bank accession no. JQ237823) were generated by transfecting MNoV cDNA clones into 293T cells as described previously [16]. For cell survival assays, 1 × 10^4^ of the indicated HeLa cells were seeded in wells of a white-walled 96-well plate (Corning) with indicated MNoV strains at an MOI of 50. 72 hours post-infection, viability was measured using CellTiter-Glo (promega) following manufactures protocol. Viability for each cell line was normalized to a mock infected sample. Each condition was performed at least in triplicate in three independent experiments. A similar experimental set up was performed for BV2 cells except an MOI of 5.0 was used and viability was assessed after 24 hours of infection.

For MNoV growth curves, 5 × 10^4^ cells infected in suspension with MNoV^CW3^ or MNoV^CR6^ at an MOI of 0.05 in a well of a 96-well plate. Plates were frozen at 12 or 24 hours post infection at −80°C for BV2 and HeLa cells, respectively. Total cell lysate was used in subsequent plaque assays as previously described [34]. All infections were done in triplicate in each of at least three independent experiments.

Viral RNA (vRNA) from MNoV^CW3^ was extracted from cell-free viral preparations using TRIzol (Invitrogen) according to manufacturer instructions. Purified vRNA was plaqued to ensure complete inactivation of MNoV. 4 μg of vRNA was transfected into BV2 cells using Lipofectamine 2000 (Invitrogen) according to the manufacturer’s protocol. Transfected cells were frozen 12 hours later. Each condition was assayed by plaque assay in duplicate in three independent experiments.

### Antibodies and Western Blotting

The following antiboides were used for flow cytometry or western blotting as indicated: rabbit α-TRIM7 (Sigma) mouse α-GAPDH-HRP (Sigma), armenian hamster α-CD300lf clone 3F6 (Genentech) anti-human CD4 (Invitrogen; clone S3.5), and antimouse CD4 (Biolegend; clone RM4-5).

Cells were placed on ice, washed once in PBS, prior to lysis in cold RIPA Buffer (50 mM Tris pH 7.4, 150 mM NaCl, 2 mM EDTA, 1% IGEPAL, 0.5% Sodium Deoxycholate, and 0.1% SDS) with HALT protease and phosphastase inhibitor cocktail (Sigma). Lysates were clarified by centrifugation prior to resolving on SDS-PAGE gels (BioRad) and transfer to PVDF membranes.

### Data availability

The data that support the findings of this study are available from the corresponding authors upon request.

## Acknowledgements

We would like to thank Megan Baldridge and Craig Wilen for helpful conversations and review of the manuscript.

---

**Figure 1:CRISPRa screen for anti-norovirus genes**

**(A)** Schematic of the CRISPRa system used in this study. Dead Cas9 (dCas9) fused to VP64 interacts with sgRNAs at the transcriptional start site of a gene. sgRNA have PP7 hairpins that recruit a chimeric protein containing Heat Shock Factor (HSF), p65 transcription factor, and phage coat protein (PCP) fused together. In combination this leads to the transcriptional activation of the targeted gene.

**(B-C)** BV2 **(B)** or HeLa-CD300lf **(C)** expressing dead Cas9-VP64 fusion were transduced with the indicated sgRNA plasmids and assayed for CD4 expression one week after antibiotic selection. At base line CD4 levels are low but dramatically increase upon expression of the CRISPRa machinery being present.

**(D)** Cartoon overview of the cell survival CRISPRa screen performed with BV2 and HeLa-CD300lf cells. sgRNAs from cells surviving MNoV challenge were analyzed and compared to relative abundance of a mock infected sample.

**Figure 2: Identification of anti-MNoV genes in mouse and human cells**

**(A)** A heat map showing enrichment of genes in the four indicated conditions. Genes are color coded based upon their STARS score.

**(B)** A comparison of the STARS score from BV2 cells challenged with MNoV^CW3^ (x-axis) and MNoV^CR6^ (y-axis). Genes that did not meet the criteria to receive a STARS score are assigned a value of 0.

**(C)** A comparison of the STARS score from HeLa-CD300lf cells challenged with MNoV^CW3^ (x-axis) and MNoV^CR6^ (y-axis). Genes that did not meet the criteria to receive a STARS score are assigned a value of 0.

**Figure 3:Validation of anti-MNoV genes in human cells**

**(A-B)** HeLa-CD300lf cells expressing dCas9-VP64 and the indicated sgRNAs were challenged with MNoVCW3 **(A)** or MNoVCR6 **(B)** at an multiplicity of infection (MOI) of 50. Cellular viability was assessed 72 hours post infection (hpi). Values are normalized for each cell line to uninfected control wells to determine % viability. Data are shown as means ± SEM from three independent experiments and data were analyzed by one-way ANOVA with Tukey’s multiple comparison test. *p<0.05, **p<0.01, ***p<0.001, ****p<0.0001, ns = not significant.

**(C)** A graph depicting the correlation between the relative viability of cell lines after 72 hours of MNoV^CW3^ (x-axis) and MNoV^CR6^ (y-axis) infection. Each dot represents the average relative viability over three independent experiments that the indicated sgRNA expressing cell line exhibited.

**(D-E)** HeLa-CD300lf cells expressing dCas9-VP64 and the indicated sgRNAs were infected with MNoVCW3 **(D)** or MNoVCR6 **(E)** at an MOI of 0.05. Viral production was assessed 24 hpi as measured by plaque assay. Data are shown as means ± SEM from three independent experiments and data were analyzed by one-way ANOVA with Tukey’s multiple comparison test. *p<0.05, **p<0.01, ***p<0.001, ****p<0.0001, ns = not significant.

**(F)** A graph depicting the correlation between the viral production 24 hours after MNoV^CW3^ (x-axis) and MNoV^CR6^ (y-axis) infection. Each dot represents the average plaque forming units (PFU)/ml over three independent experiments that the indicated sgRNA expressing cell line exhibited.

**Figure 4: A specific TRIM7 isoform inhibits MNoV replication post-entry**

**(A)** (Top) Cartoon diagram of TRIM7 isoform 1 and isoform 4. RING, B-box, coiled-coil (CC) and SPRY domains are displayed. Isoform 4 has an alternative, shorter coiled coil domain and does not encode a SPRY domain. (Bottom) Representative western blot for indicated protein expression in BV2 or HeLa-CD300lf cells expressing either an empty vector, TRIM7 isoform 1 or TRIM7 isoform 4.

**(B-C)** HeLa-CD300lf cells expressing indicated constructs infected an multiplicity of infection (MOI) of 50 with MNoV^CW3^ **(B)** or MNoV^CR6^ **(C)**. Cellular viability was assessed 72 hours post infection (hpi). Values are normalized for each cell line to uninfected control wells to determine % viability. Data are shown as means ± SEM from three independent experiments and data were analyzed by one-way ANOVA with Tukey’s multiple comparison test **p<0.01, ***p<0.001, ns = not significant.

**(D-E)** BV2 cells expressing indicated constructs infected an multiplicity of infection (MOI) of 5 with MNoV^CW3^ (D) or MNoV^CR6^ (E). Cellular viability was assessed 24 hours post infection (hpi). Values are normalized for each cell line to uninfected control wells to determine % viability. Data are shown as means ± SEM from three independent experiments and data were analyzed by one-way ANOVA with Tukey’s multiple comparison test ***p<0.001, ****p<0.0001, ns = not significant.

**(F-G)** HeLa-CD300lf cells expressing indicated constructs infected an multiplicity of infection (MOI) of 0.05 with MNoV^CW3^ (F) or MNoV^CR6^ **(G)**. Viral production was assessed 24 hpi as measured by plaque assay. Data are shown as means ± SEM from three independent experiments and data were analyzed by one-way ANOVA with Tukey’s multiple comparison test. ****p<0.0001, ns = not significant.

**(H-I)** BV2 cells expressing indicated constructs infected an multiplicity of infection (MOI) of 0.05 with MNoV^CW3^ **(H)** or MNoV^CR6^ **(I)**. Viral production was assessed 12 hpi as measured by plaque assay. Data are shown as means ± SEM from three independent experiments and data were analyzed by one-way ANOVA with Tukey’s multiple comparison test. ****p<0.0001, ns = not significant.

**(J)** BV2 expressing indicated constructs were transfected with viral RNA from MNoV^CW3^ and harvested 12 hours post-transfection. Viral production was measured by plaque assay. Data are shown as means ± SEM from three independent experiments and data were analyzed by one-way ANOVA with Tukey’s multiple comparison test. ****p<0.0001, ns = not significant.

## Supporting Information Legends

**Supplemental Methods:** DNA sequences used for this study

**Table S1:** STARS analysis of CRISPRa screen in BV2 cells for MNoV^CW3^

**Table S2:** STARS analysis of CRISPRa screen in BV2 cells for MNoV^CR6^

**Table S3:** STARS analysis of CRISPRa screen in HeLa-CD300lf cells for MNoV^CW3^

**Table S4:** STARS analysis of CRISPRa screen in HeLa-CD300lf cells for MNoV^CR6^

